# Grazing cattle respond more strongly to a quadruped robot than to a drone but habituate over consecutive days

**DOI:** 10.64898/2026.08.12.744347

**Authors:** Hiroki Anzai, Kirika Iwatani, Sakuya Miyamoto, Kanon Tamura, Honoka Miyagi

## Abstract

Robots are increasingly used to manage animals, but whether animals keep responding to robots over repeated encounters is poorly understood. Habituation theory predicts that more salient stimuli elicit stronger responses and habituate more slowly, and that responses recover partially after a pause in stimulation, followed by faster habituation. We tested these predictions in a field study of robotic herding of grazing Japanese Black cattle, *Bos taurus*. We compared the responses of 22 cows to a drone and to a quadruped robot, each playing either dog barking or an air horn (266 approach events), and herded 24 cows with the robot on five consecutive days in September and three consecutive days in October, after a 24-day interruption (823 approach events). Binomial generalized linear mixed models with cow identity as a random effect showed that the odds of moving away from the device were twice as high for the robot as for the drone, the odds of showing no response were six times lower and the odds of flight were four times higher; the acoustic stimulus had no main effect on any response. Herding succeeded more often with the robot, particularly when it played dog barking. During consecutive herding with the robot, the odds of moving away and of flight declined by approximately 30% per day in September, responses partially recovered after the interruption, and the decline in October was steeper than in September for flight. Fear-related responses occurred in fewer than 5% of approach events, and cows differed consistently in their responsiveness. Our results show that a ground-level, dog-like robot is a more salient stimulus for cattle than a drone, but its effect still wanes with repeated exposure, in line with the characteristics of habituation described in laboratory studies. Managing habituation will therefore be essential for sustained robotic herding.

---

Animals increasingly encounter robots in agricultural and natural environments, and robots are also used as experimental tools to probe animal behaviour (Papadopoulou et al., 2025). Whether animals continue to respond to a robot over repeated encounters is central to both uses: an animal that rapidly ceases to respond renders a robot ineffective as a management tool, and changes in responsiveness alter what a robot measures when it is used as an experimental stimulus. Robotic herding, in which robots direct the movement of animals, provides a tractable system for studying these dynamics under natural conditions (King et al., 2023; Papadopoulou et al., 2025).

Habituation, the decline in response to a repeated stimulus that is not explained by sensory adaptation or fatigue, shows several parametric characteristics (Rankin et al., 2009). Among them, more intense stimuli produce slower and less pronounced habituation; responses recover at least partially when the stimulus is withheld (spontaneous recovery); and after repeated series of stimulation and recovery, habituation becomes progressively faster (potentiation of habituation). These characteristics were established largely in laboratory studies with short interstimulus intervals, and it is less clear whether they hold when free-ranging animals encounter a moving agent once a day over days or weeks.

Because grazing animals use pasture selectively, their spatial distribution tends to be uneven, causing local overgrazing, soil erosion and inefficient forage use (Distel et al., 2004). Robotic herding could improve grazing distribution with little labour, and a ground vehicle elicited avoidance in cattle with few signs of fear (Anzai & Sakurai, 2022). However, cattle rapidly habituated to daily herding by a drone, and their responsiveness did not recover after a 24-day interruption (Anzai & Kumaishi, 2023). A drone flying above grazing animals may be a weak stimulus because it stays largely outside their horizontal field of view, which, according to habituation theory, would explain the rapid habituation.

A quadruped robot moves on the ground at the eye level of grazing animals and resembles a dog in body form and gait. Sheep in an exploratory trial appeared to respond to a quadruped robot much as they respond to a sheepdog (Ravikanna et al., 2025), dogs distinguished a dog-like robot from a live conspecific but still showed social responses towards it (Kubinyi et al., 2004), and wild vervet monkeys showed alarm responses on first encountering a quadruped robot but habituated within days (Canteloup et al., 2024). No study has yet quantified how livestock respond to a quadruped robot or whether such responses persist under repeated exposure.

We studied the responses of grazing cattle to robotic herding in two phases. In comparison trials, cows were approached by either a drone or a quadruped robot. In continuous trials, cows were herded with the robot on consecutive days before and after a 24-day interruption. We tested three predictions derived from habituation theory. (1) If ground-level approach and a dog-like form make a quadruped robot a more salient stimulus than a drone, cows should respond more strongly to the robot (more frequent moving away and flight, shorter latency and fewer non-responses), and herding should succeed more often with the robot. (2) Repeated daily herding with the robot should lead to a progressive decline in responsiveness. (3) After the 24-day interruption, responsiveness should partially recover, and the subsequent decline should be faster than during the first exposure period. Because the robot played one of two sounds, we also examined whether the acoustic stimulus modified the difference between devices.

## METHODS

### Study Site and Animals

The study was conducted from May to October 2023 at the Sumiyoshi Livestock Science Station of the University of Miyazaki (31°59′N, 131°28′E), southern Kyushu, Japan. The experimental pasture was a relatively flat 1.1 ha sward dominated by centipedegrass, *Eremochloa ophiuroides*, and bahiagrass, *Paspalum notatum*, with a low cover of Japanese lawngrass, *Zoysia japonica*. The pasture was divided into nine equal plots arranged in a 3 × 3 grid (A1–C3), marked by poles (1.5 m high, with yellow markers) at the corners of each plot (Fig. 1). A 0.4 ha resting area with water and shade trees adjoined the pasture and was accessible through a single gate in plot A1; cattle could move freely between the pasture and the resting area at any time during grazing hours. A separate gate between the resting area and the barn was closed during grazing hours.

**Figure 1.**
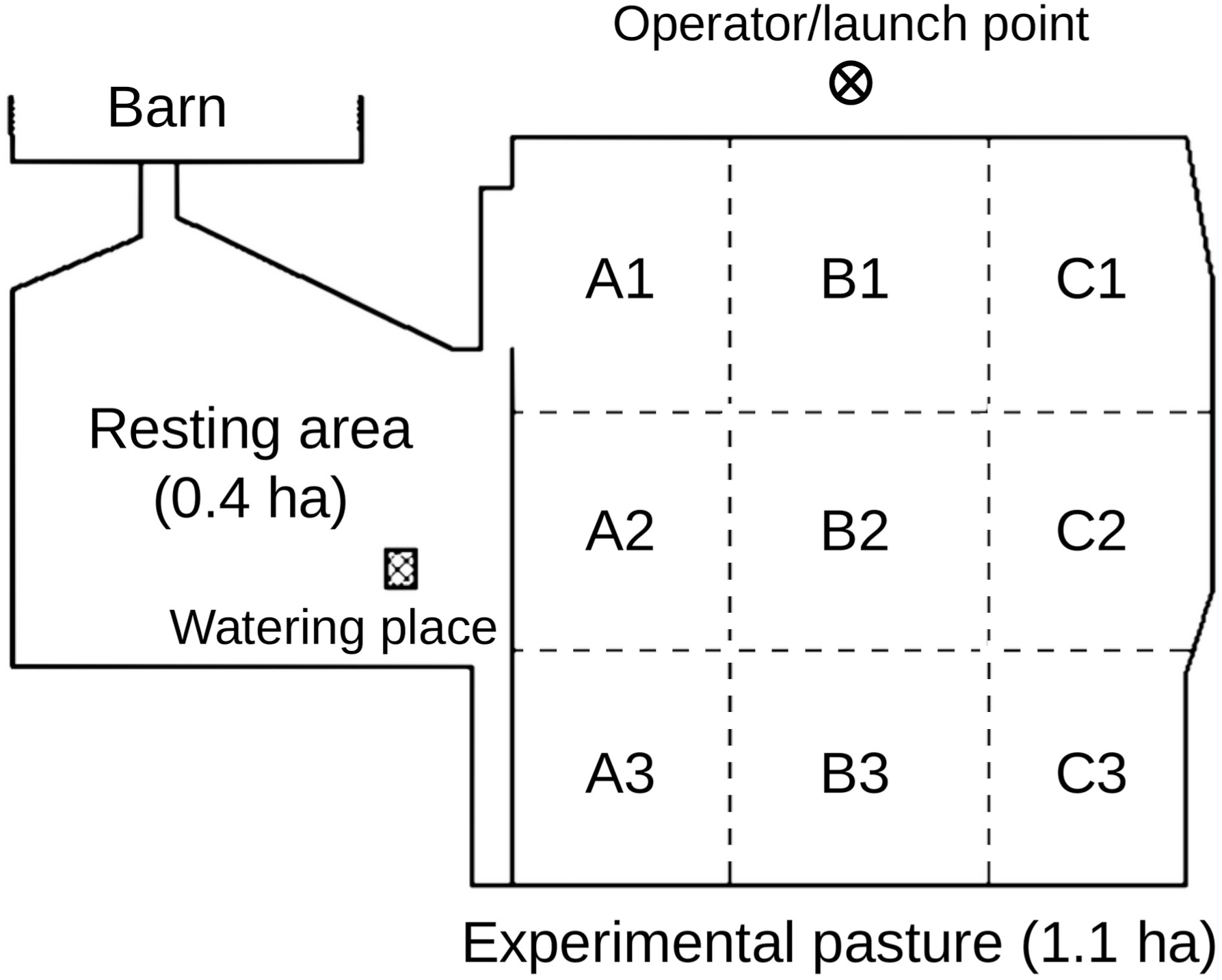
Layout of the experimental pasture. A1–C3 denote nine equal plots in a 3 × 3 grid. The herding area comprised plots B1 and C1 in the comparison trials and plots A1, B1 and C1 in the continuous trials.

A herd of 25–29 Japanese Black breeding cows (all female; mean ± SE age = 10.2 ± 0.7 years) and seven to 11 calves grazed the experimental pasture for five consecutive days each month from May to October (0900–1600 hours each day). Outside grazing hours and during the nongrazing season (November–April), all animals were housed in a barn and were therefore thoroughly accustomed to humans. They had no direct experience with herding dogs but occasionally heard dogs from adjacent areas. Herd composition was largely stable, although a few individuals entered or left the herd because of parturition or other management reasons. The cows included in each analysis are described in Statistical Analysis.

### Herding Devices

We used two herding devices: a drone (Mavic 2 Enterprise DUAL; DJI Technology, Shenzhen, Guangdong, China) and a quadruped robot (Unitree Go1; Unitree Robotics, Hangzhou, Zhejiang, China) (Fig. 2). The drone measured 322 × 242 × 140 mm (length × width × height), weighed 899 g at take-off and was grey; its maximum flight time was 31 min and its maximum horizontal speed was approximately 14 m/s. The robot measured 640 × 280 × 400 mm, weighed 13 kg including the battery and was black; its walking speed on the pasture was approximately 0.9 m/s. The robot could move forwards, backwards and sideways, turn and move its head up and down.

**Figure 2.**
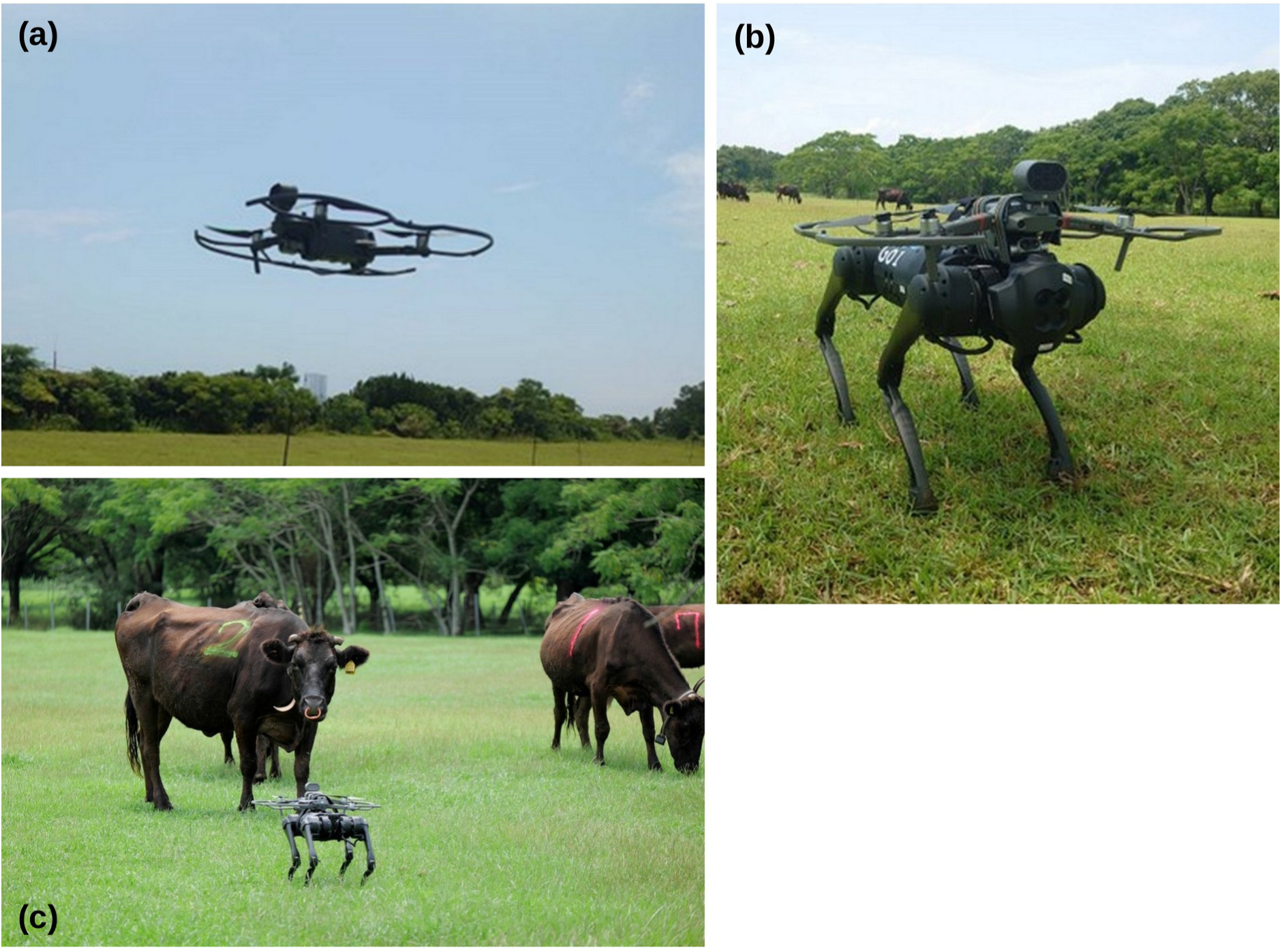
Herding devices. (a) Drone (Mavic 2 Enterprise DUAL). (b) Quadruped robot (Unitree Go1) with the drone attached to its back to deliver the acoustic stimulus. (c) The quadruped robot approaching a cow during a herding trial.

Sounds were played through the speaker accessory of the drone (maximum output 100 dB at 1 m), which was mounted on the drone. Sounds were played continuously during each approach. When the robot was used, the drone was attached to the back of the robot; in this configuration only the speaker was used, and the propellers and all other systems were inactive.

### Herding Procedure

Herding trials were conducted during grazing hours on 13 days between June and October. Because continuous operation was limited by battery charging, it was not feasible to exclude cows completely from heavily grazed areas. We therefore used the devices to drive cows out of heavily grazed areas and to discourage their use of these areas. We refer to the area from which cows were driven out as the herding area. The herding area for each month was selected on the basis of the vegetation before grazing.

A herding session was started whenever two or more cows were grazing within the herding area. The operator manoeuvred the drone or robot towards a target cow from any direction except the rear blind area. The drone was flown at a height of 1–2 m, approximately at the eye level of grazing cattle. If a cow did not move away when the device came within approximately one cow length, the drone was moved sideways, or the robot moved sideways, turned and moved its head up and down in random order for approximately 20 s. If the cow remained unresponsive and other cows were grazing in the herding area, the operator switched to another cow. A session ended when all cows had left the herding area or when all targeted cows had remained unresponsive for approximately 15–20 min. In the latter case, a new session was started after a further 15–20 min if cows were still in the herding area. Sessions were also interrupted when the robot overheated or needed a battery change. Calves were not targeted. A single operator operated the devices throughout the study and remained outside the herding area, concealed behind vegetation.

### Experimental Design

The comparison trials compared the drone and the quadruped robot. The two devices were alternated between herding sessions. Trials were conducted on the last grazing day of June and July and on the first three grazing days of August (five days in total). The herding area comprised plots B1 and C1 (approximately 22% of the pasture area). Dog barking and an air horn were used as acoustic stimuli with both devices. Dog barking was selected for the continuous trials because it has been reported to be an effective directional cue in sheep herding (Yaxley et al., 2021) and because a preliminary analysis indicated no difference between the two sounds.

The continuous trials examined daily changes in responses during consecutive herding and whether responses recovered after a 24-day interruption. Trials were conducted on all five grazing days in September (16–20 September; Days 1–5) and the first three grazing days in October (15–17 October; Days 6–8). Only the quadruped robot was used, with dog barking. The herding area was expanded to three plots (A1, B1 and C1; approximately 33% of the pasture area) to collect more data.

Trial days were mostly sunny. Maximum air temperatures exceeded 30 °C on the trial days in July–September. High temperatures occasionally caused the robot to overheat, so that herding could not be conducted even when cattle were in the herding area. On Day 3 of the continuous trials, light rain fell during grazing hours, and herding was interrupted for approximately 30 min. Mean wind speeds were low throughout and did not affect the operation of the drone.

### Behavioural Observations

All herding trials were video-recorded (HDR-CX180; Sony, Tokyo, Japan). Four responses were scored for each approach event: (1) type of first behavioural response, (2) latency to first response, (3) occurrence of flight and (4) success or failure of herding.

The type of first response was classified into five categories according to the ethogram in Table 1. Latency to first response was the time from when the device came within one cow length of the target cow to when the cow stopped grazing; if the cow began responding before the device came within one cow length, latency was recorded as 0 s. Latency was classified into four categories: 0 s, 1–9 s, 10–19 s and ≥20 s; cows that did not respond were assigned to ≥20 s. Flight was recorded when the cow moved more than one cow length away from the approaching device, regardless of the type of first response. Herding was recorded as successful if the cow left the herding area during the session, regardless of the direct cause. When the same cow was approached more than once within a session, only the first approach was analysed for the type of first response, latency and flight, whereas herding success was the final outcome for that cow in the session. Outcomes that could not be confirmed from the video or field records were classified as unknown and excluded from the analysis of herding success.

**Table 1.** Ethogram used to classify the first behavioural response of cows to the approaching device.

| Behaviour | Description |
| --- | --- |
| Move away | Walking $\geq 5$ steps away from the approaching device |
| Interest | Approaching the device, or staring at it for $\geq 10$ s |
| Fear/startle/attack | Jumping, running, backing, attacking or trying to attack in response to the approaching device |
| Slight reaction | Moving $\leq 4$ steps while continuing to graze, or briefly orienting towards the device before returning to grazing |
| No response | Continuing to graze without any detectable reaction to the approaching device |

Calves were present in the herding area but were not targeted. Their responses to the robot during the continuous trials are described in the Supplementary Material.

### Statistical Analysis

We analysed five response variables for each approach event. Four were binary: whether the first response was moving away, whether the cow showed no response, whether flight occurred and whether herding succeeded. Of the five types of first response, we analysed moving away and no response because they represent the two ends of the response to herding (the device was effective or had no effect); interest and fear/startle/attack were too rare for separate models (about 5% and 3% of events), and slight reaction is intermediate, so these three types are reported descriptively. The binary variables were analysed with generalized linear mixed models (GLMMs) with a binomial error distribution, a logit link and cow identity as a random intercept, to account for repeated observations of the same cows (Bolker et al., 2009). Latency to first response was analysed as an ordinal variable with four categories (0 s, 1–9 s, 10–19 s and ≥20 s) in a cumulative logit GLMM with the same random effect; for latency, an odds ratio above 1 indicates shorter latency. Models were fitted by maximum likelihood with the Laplace approximation in PROC GLIMMIX of SAS 9.4 (SAS Institute, Cary, NC, U.S.A.), with denominator degrees of freedom calculated by the between–within method. Fixed effects were tested with *F* tests, and we report odds ratios (OR) with 95% confidence intervals (CI) as effect sizes. All tests were two tailed, with *α* = 0.05.

The comparison trials included the 22 cows that remained in the herd throughout June–August (266 approach events; 234 events with a known herding outcome). Each model included device (drone or robot) and acoustic stimulus (dog barking or air horn) as fixed effects. To test whether the difference between devices depended on the sound, we fitted a second model that added the device × stimulus interaction. When the interaction was significant, we estimated the effect of device separately for each sound.

The continuous trials included the 24 cows that remained in the herd throughout September and October (823 approach events; 516 events with a known herding outcome); 22 of these cows had taken part in the comparison trials. To test for recovery after the interruption (prediction 3), we fitted models with trial day as a categorical factor (eight levels) and compared Day 6 with Day 5 (recovery) and Day 6 with Day 1 (completeness of recovery) using planned contrasts. To test for a progressive decline (prediction 2) and for faster decline after the interruption (prediction 3), we fitted models with month (September or October), day within month (a covariate: 1–5 in September and 1–3 in October) and their interaction. From these models we estimated the change in the odds of each response per day within each month and tested whether the two rates differed. Key results are summarized in Tables 2–4, and full model estimates are given in Tables S1 and S2.

**Table 2.** Effects of device and acoustic stimulus on the responses of cows in the comparison trials.

| Response | Drone, %<br>(*N*) | Robot, % (*N*) | Device: OR (95%<br>CI) | Device | Sound | Device × sound |
| --- | --- | --- | --- | --- | --- | --- |
| Move away | 42.6 (115) | 58.9 (151) | 2.02 (1.21–3.37) | $F_{1,242} = 7.31, P = 0.007$ | $F_{1,242} = 0.02, P = 0.876$ | $F_{1,241} = 0.02, P = 0.890$ |
| No response | 22.6 (115) | 4.6 (151) | 0.16 (0.06–0.39) | $F_{1,242} = 15.94, P < 0.001$ | $F_{1,242} = 0.16, P = 0.688$ | $F_{1,241} = 0.79, P = 0.375$ |
| Latency<sup>a</sup> | 46.1 (115) | 60.3 (151) | 2.58 (1.56–4.28) | $F_{1,242} = 13.69, P < 0.001$ | $F_{1,242} = 0.56, P = 0.456$ | $F_{1,241} = 4.43, P = 0.036$ |
| Flight | 52.2 (115) | 82.1 (151) | 4.18 (2.38–7.34) | $F_{1,242} = 24.95, P < 0.001$ | $F_{1,242} = 0.19, P = 0.665$ | $F_{1,241} = 0.13, P = 0.718$ |
| Herding success<sup>b</sup> | 60.8 (97) | 84.7 (137) | 4.30 (2.15–8.59) | $F_{1,210} = 17.25, P < 0.001$ | $F_{1,210} = 0.11, P = 0.744$ | $F_{1,209} = 4.20, P = 0.042$ |

Percentages are proportions of approach events (22 cows). Device and sound effects are from binomial GLMMs (cumulative logit GLMM for latency) with cow identity as a random intercept; the interaction was tested in a separate model. Odds ratios compare the robot with the drone. Full estimates are given in Tables S1 and S2.
<sup>a</sup> Percentages are events with a latency of 0 s; the model used all four latency categories, and an odds ratio above 1 indicates shorter latency with the robot.
<sup>b</sup> Events with a known outcome.

**Table 3.** Changes across trial days and recovery after the 24-day interruption in the continuous trials.

| Response | Trial day | Day 6 vs Day 5: OR (95% CI) | Day 6 vs Day 1: OR (95% CI) | Cow variance |
| --- | --- | --- | --- | --- |
| Move away | $F_{7,155} = 7.38, P < 0.001$ | 1.82 (1.05–3.16), $P = 0.034$ | 0.52 (0.26–1.04), $P = 0.065$ | $0.73 \pm 0.27$ |
| No response | $F_{7,155} = 7.11, P < 0.001$ | 0.26 (0.11–0.62), $P = 0.003$ | 2.85 (0.70–11.56), $P = 0.142$ | $4.41 \pm 1.87$ |
| Latency<sup>a</sup> | $F_{7,155} = 5.32, P < 0.001$ | 1.86 (1.10–3.15), $P = 0.020$ | 0.43 (0.21–0.86), $P = 0.018$ | $1.84 \pm 0.60$ |
| Flight | $F_{7,155} = 8.97, P < 0.001$ | 1.77 (0.93–3.38), $P = 0.080$ | 0.37 (0.15–0.93), $P = 0.035$ | $2.00 \pm 0.73$ |
| Herding success<sup>b</sup> | $F_{7,141} = 3.73, P = 0.001$ | 0.58 (0.24–1.42), $P = 0.232$ | 0.56 (0.20–1.55), $P = 0.261$ | $0.31 \pm 0.21$ |

GLMMs with trial day (Days 1–8) as a categorical fixed effect and cow identity as a random intercept (24 cows; 823 approach events). Contrasts are planned comparisons ( $t$ tests with 155 df, or 141 df for herding success). Cow variance is the estimated variance of the random intercept ± SE.
<sup>a</sup> Cumulative logit GLMM; an odds ratio above 1 indicates shorter latency.
<sup>b</sup> 516 events with a known outcome.

**Table 4.** Daily rate of change in the responses of cows within September and October in the continuous trials.

| Response | September: OR per day (95% CI) | October: OR per day (95% CI) | Ratio October/September (95% CI) | Month × day |
| --- | --- | --- | --- | --- |
| Move away | 0.72 (0.62–0.84) | 0.55 (0.41–0.73) | 0.76 (0.55–1.05) | $F_{1,796} = 2.86, P = 0.091$ |
| No response | 1.85 (1.40–2.44) | 2.70 (1.78–4.11) | 1.46 (0.90–2.39) | $F_{1,796} = 2.34, P = 0.127$ |
| Latency<sup>a</sup> | 0.69 (0.60–0.81) | 0.71 (0.55–0.92) | 1.02 (0.77–1.37) | $F_{1,796} = 0.03, P = 0.870$ |
| Flight | 0.67 (0.55–0.81) | 0.44 (0.32–0.61) | 0.66 (0.46–0.95) | $F_{1,796} = 5.08, P = 0.025$ |
| Herding success<sup>b</sup> | 0.86 (0.67–1.12) | 0.87 (0.58–1.32) | 1.01 (0.62–1.64) | $F_{1,489} = 0.00, P = 0.974$ |

GLMMs with month (September or October), day within month (covariate; Days 1–5 in September and Days 6–8 in October) and their interaction as fixed effects and cow identity as a random intercept (24 cows). The odds ratio per day describes the daily change in the odds of each response; the month × day interaction tests whether the rate differed between months.
<sup>a</sup> Cumulative logit GLMM; an odds ratio below 1 indicates that latency became longer.
<sup>b</sup> 516 events with a known outcome.

### Ethical Note

All procedures were approved by the Animal Care and Use Committee of the University of Miyazaki (approval number 2020-006-5) and followed the ASAB/ABS Guidelines for the ethical treatment of nonhuman animals and the ARRIVE guidelines. The cows belonged to the research herd of the station and were kept under its routine management throughout; no animals were purchased, transported or removed from the herd for this study. The number of cows was determined by the size of the herd: we included all cows present throughout each trial period, and no a priori power analysis was possible. During grazing hours, cows had free access to water and shade in the resting area; outside grazing hours they were housed in a barn, without bedding, and were fed silage and hay in appropriate amounts with water available ad libitum.

The approach of a dog-like robot and the playback of dog barking could be perceived as predator-like stimuli and might cause stress. To minimize this, the devices never touched the animals, approaches were not pursued closer than approximately one cow length, cows were never confined and could always move away from the device or into the resting area, and sessions ended when targeted cows remained unresponsive for 15–20 min. Calves were never targeted. Herding was suspended during rain. Some cows were pregnant during the trials; because the devices never made contact and cows could always move away at their own pace, we considered the risk to pregnant cows to be low. Under the approved protocol, a trial was to be stopped if any animal showed strong signs of stress, but no such case occurred. Fear-related responses (jumping, running, backing or attacking) occurred in fewer than 5% of approach events, only one attack on the robot was recorded during the continuous trials, and no animal was harmed. At the end of the study, all animals remained in the station herd under routine management.

## RESULTS

### Device Comparison

In the comparison trials, we recorded 266 approach events for 22 cows (drone: *N* = 115; robot: *N* = 151; median = 11.5 events per cow, range 4–25). The robot elicited stronger responses than the drone in all five response variables (Table 2, Fig. 3). Cows were more likely to move away from the robot (58.9% versus 42.6% of events; OR = 2.02, 95% CI = 1.21–3.37) and less likely to show no response (4.6% versus 22.6%; OR = 0.16, 95% CI = 0.06–0.39). They responded sooner to the robot (latency of 0 s: 60.3% versus 46.1%; ≥20 s: 9.9% versus 33.0%; OR = 2.58, 95% CI = 1.56–4.28) and fled more often (82.1% versus 52.2%; OR = 4.18, 95% CI = 2.38–7.34). Herding succeeded more often with the robot (84.7% versus 60.8% of events with a known outcome; OR = 4.30, 95% CI = 2.15–8.59).

**Figure 3.**
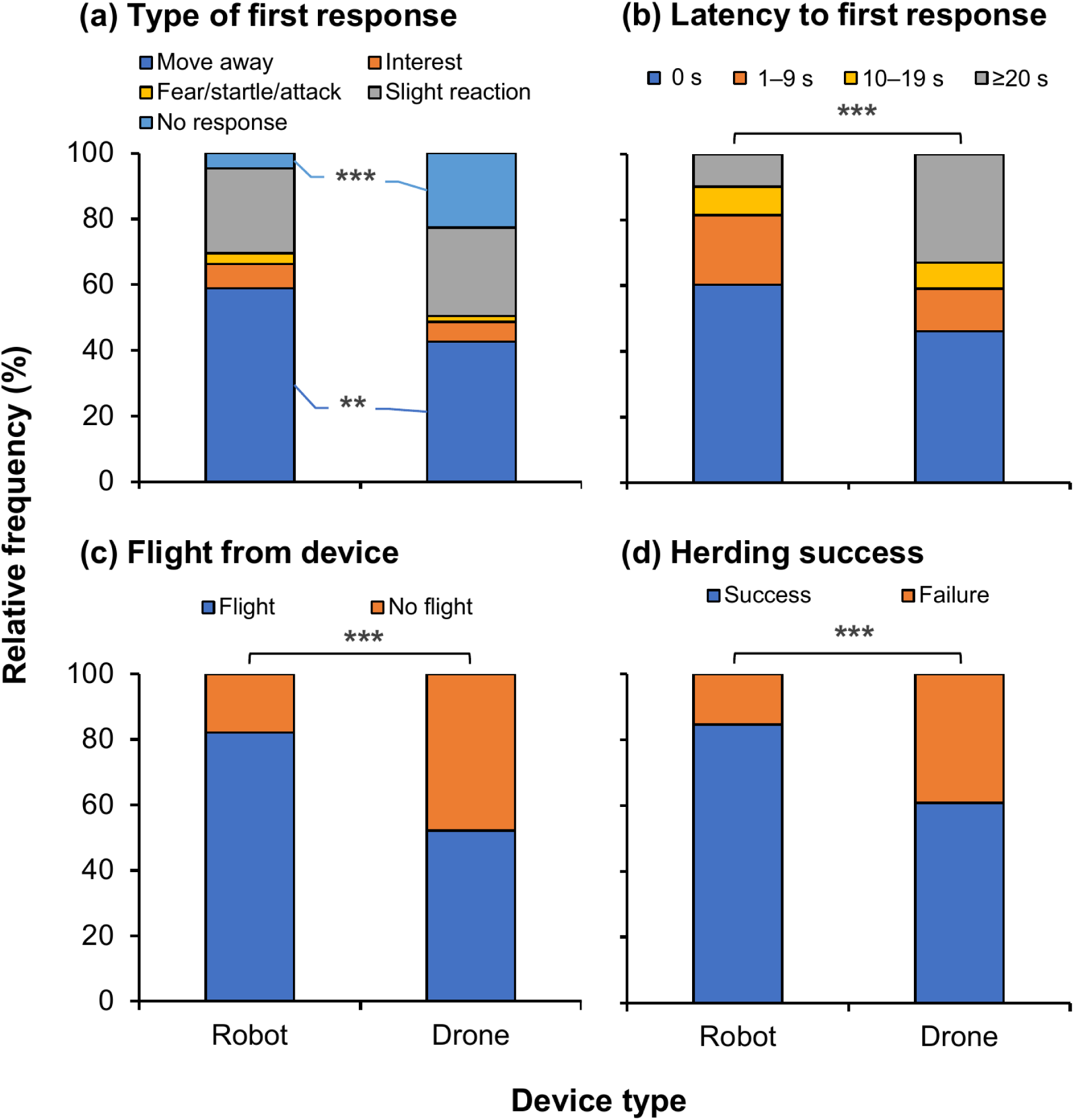
Responses of cows to the drone and the quadruped robot in the comparison trials (22 cows). Percentage of approach events by (a) type of first response, (b) latency to first response, (c) occurrence of flight and (d) herding success. Drone: *N* = 115 events; robot: *N* = 151 events. Herding success was calculated for events with a known outcome (drone: *N* = 97; robot: *N* = 137). Asterisks indicate differences between devices in GLMMs (Table 2): \**P* < 0.05; \*\**P* < 0.01; \*\*\**P* < 0.001.

The acoustic stimulus had no main effect on any response, and it did not modify the difference between devices for moving away, no response or flight (Table 2). For herding success and latency, however, the difference between devices depended on the sound (Table 2). With dog barking, herding succeeded more often with the robot than with the drone (88.7%, *N* = 71, versus 50.0%, *N* = 38; OR = 9.13, 95% CI = 3.23–25.84, *P* < 0.001), whereas with the air horn the difference was smaller and not significant (80.3%, *N* = 66, versus 67.8%, *N* = 59; OR = 2.30, 95% CI = 0.95–5.58, *P* = 0.065). For latency, the pattern was reversed: the robot shortened latency with the air horn (0 s latency: 61.5% versus 38.2%; OR = 4.09, 95% CI = 2.09–8.02, *P* < 0.001) but not with dog barking (58.9% versus 57.4%; OR = 1.39, 95% CI = 0.65–2.99, *P* = 0.40), because cows already responded quickly to the drone when it played dog barking (dog barking versus air horn with the drone: OR = 2.25, 95% CI = 1.04–4.88, *P* = 0.039).

### Daily Changes During Continuous Herding

In the continuous trials, we recorded 823 approach events for 24 cows (59–136 events per day; median = 32 events per cow, range 13–55). Across all days, the first response was moving away in 43.0% of events, slight reaction in 32.8%, no response in 16.0%, interest in 5.1% and fear/startle/attack in 3.0%; only one attack was recorded. All five response variables changed across trial days (Table 3, Fig. 4).

**Figure 4.**
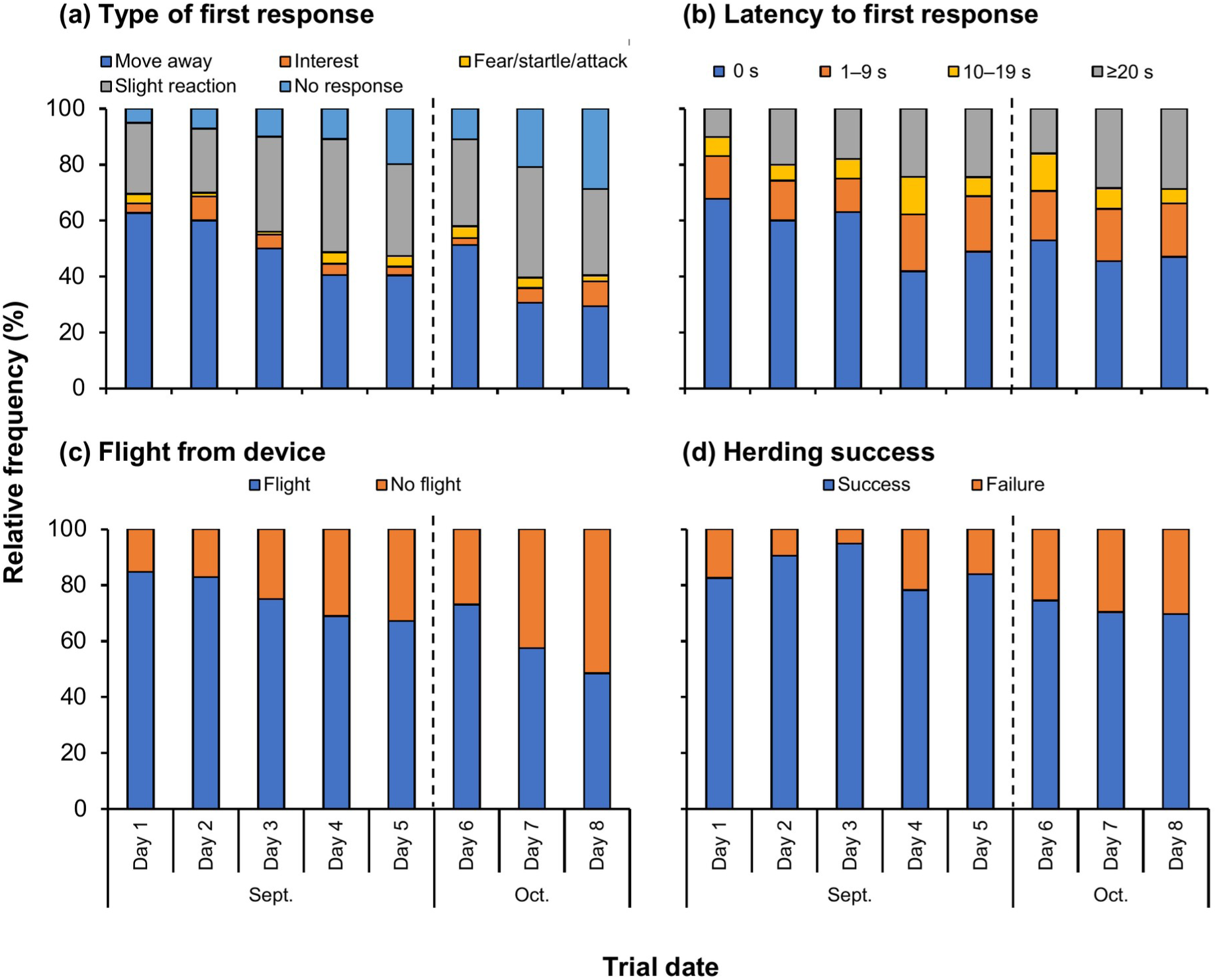
Daily changes in the responses of cows to the quadruped robot in the continuous trials (24 cows). Percentage of approach events by (a) type of first response, (b) latency to first response, (c) occurrence of flight and (d) herding success on each trial day. Days 1–5: 16–20 September; Days 6–8: 15–17 October. The dashed vertical line indicates the 24-day interruption. Statistical tests are given in Tables 3 and 4.

In September, responsiveness declined day by day (Table 4). The odds of moving away decreased by 28% per day (from 62.7% of events on Day 1 to 40.5% on Day 5), the odds of flight by 33% per day (from 84.7% to 67.2%), and latency became longer (0 s latency: from 67.8% to 48.9%), whereas the odds of no response increased by 85% per day (from 5.1% to 19.8%).

After the interruption, responsiveness partially recovered (Table 3). Compared with Day 5, cows on Day 6 were more likely to move away (51.3% versus 40.5%; OR = 1.82, 95% CI = 1.05–3.16), less likely to show no response (10.9% versus 19.8%; OR = 0.26, 95% CI = 0.11–0.62) and responded sooner (OR = 1.86, 95% CI = 1.10–3.15); flight showed a similar but nonsignificant increase (73.1% versus 67.2%; *P* = 0.080). Recovery was incomplete: flight and short latencies remained less frequent on Day 6 than on Day 1 (*P* = 0.035 and *P* = 0.018), and moving away tended to be less frequent (*P* = 0.065).

In October, responsiveness declined again (Table 4). Moving away fell from 51.3% on Day 6 to 29.4% on Day 8 (OR per day = 0.55) and flight from 73.1% to 48.5% (OR per day = 0.44), whereas no response rose from 10.9% to 28.7%. The daily decline in flight was steeper in October than in September (*P* = 0.025). Moving away and no response showed the same tendency without reaching significance (*P* = 0.091 and *P* = 0.13), and the rate of change in latency did not differ between months (*P* = 0.87).

Herding success was 81.4% overall (516 events with a known outcome). It was highest on Day 3 (94.9%) and lower on Days 6–8 (69.7–74.5%) than on any day in September (78.3–94.9%), but it showed no consistent daily trend within either month and did not differ between Day 6 and Day 5 (Tables 3, 4).

Cows differed consistently in their responsiveness: the variance of the random intercept for cow identity was large for no response, latency and flight (Table 3).

## DISCUSSION

Our results supported the prediction that cattle respond more strongly to a quadruped robot than to a drone (prediction 1). Cows approached by the robot were more likely to move away and to flee, responded with shorter latency and rarely ignored the device, and herding succeeded more often. These differences remained after accounting for the acoustic stimulus and for repeated observations of the same cows. The robot elicited stronger responses even though the drone was flown at 1–2 m, approximately at the eye level of grazing cattle. The two devices differed not only in their position relative to the cattle and their movement but also in body size, colour and gait, and the present data cannot separate these factors. The quadrupedal gait and body form of the robot may have resembled a herding dog or a predator more closely than the aerial movement of a drone. Experiments that vary these features systematically are needed to identify which features of quadruped robots drive the responses of livestock.

The acoustic stimulus modified the difference between devices for two responses, but in opposite directions. The advantage of the robot for herding success was largest when it played dog barking. A dog-like robot that also sounds like a dog presents congruent visual and acoustic cues, and congruent multisensory signals can elicit stronger responses than either component alone, as shown in squirrels responding to a robotic conspecific (Partan et al., 2009). For latency, in contrast, the advantage of the robot was smaller with dog barking, because cows already responded quickly to the drone when it played dog barking. These interactions were not anticipated and were not consistent across responses, so they should be regarded as exploratory and tested in experiments designed for this purpose.

As predicted (prediction 2), responsiveness to the robot declined progressively over the five consecutive days in September: cows became less likely to move away and to flee, and more likely to continue grazing without any detectable reaction. This pattern is consistent with habituation to repeated stimulation (Rankin et al., 2009) and parallels the decline reported for drone herding of cattle (Anzai & Kumaishi, 2023). The decline appeared slower than with the drone, however: in the present study, the proportion of no response increased from 5.1% to 28.7% over the eight trial days, compared with an increase from 40.6% to 78.4% over eight days with the drone (Anzai & Kumaishi, 2023). This comparison across studies is consistent with the prediction that a more salient stimulus habituates more slowly, but it must be interpreted with caution: the two studies differed in acoustic stimulation, season, sward condition, temperature and size of the herding area, and the cows in the present study had already encountered both devices during the comparison trials. Behavioural adjustments to a novel robot have also been documented in housed cattle: dairy cows reduced the time spent standing in walkways used by a robotic scraper and increased lying and feeding time (Doerfler et al., 2016).

Consistent with prediction 3, responsiveness partially recovered after the 24-day interruption: cows moved away more often, ignored the robot less often and responded sooner on Day 6 than on Day 5, although flight and short latencies remained less frequent than on Day 1. This partial recovery contrasts with the absence of recovery after the same interval with a drone (Anzai & Kumaishi, 2023) and suggests that the stronger stimulation by the robot maintained residual wariness. Responsiveness then declined again over Days 7 and 8; for flight the decline was significantly steeper than in September, and moving away and no response showed the same, nonsignificant tendency. On Day 8, interest in the robot reached its highest level (8.8% of events), suggesting a shift from avoidance towards investigation. Faster habituation after a period of recovery is the pattern described as potentiation of habituation (Rankin et al., 2009). Our results suggest that spontaneous recovery and potentiation, characteristics established mainly with short interstimulus intervals in laboratory preparations, also describe the responses of free-ranging animals to an agent encountered once a day over several weeks. Similar habituation within days has been reported for wild vervet monkeys encountering a quadruped robot (Canteloup et al., 2024), and limited behavioural responses to drone flights despite marked physiological responses have been reported in American black bears (Ditmer et al., 2015).

Cows differed markedly and consistently in their responsiveness to the robot, as shown by the large between-cow variance in no response, latency and flight. Such consistent individual differences in reactions to novel or threatening stimuli are a general feature of animal behaviour (Réale et al., 2007) and may shape collective outcomes during herding, because the movement of responsive individuals can draw less responsive group members along. This may explain why herding success, unlike the responses of individual cows, did not decline steadily within each month and was highest on Day 3, when some cows had already begun to habituate: cows that moved away may have been followed by less responsive herd mates. For robotic herding, this suggests that targeting responsive individuals could be an efficient strategy, a possibility that requires data on individual roles in collective movement.

The present study demonstrates that a quadruped robot can be used to herd grazing cattle with few signs of fear. Fear-related responses were rare with both devices, indicating that neither the devices nor the sounds caused strong aversion. This is an important prerequisite for practical use, because excessively aversive stimuli would compromise animal welfare (Herlin et al., 2021). The main obstacle to practical use remains habituation. To maintain effective herding over consecutive days, stimuli must be aversive enough to slow habituation while remaining acceptable for animal welfare. Combining sensory modalities is one promising approach, because multisensory stimulation can enhance responses beyond those elicited by either modality alone (Partan et al., 2009). Rotating among different stimuli may also slow habituation by reducing the predictability of the stimulus. More broadly, the choice of robotic agent, the intensity and variety of stimuli and the scheduling of deployment must be considered together to sustain the influence of robots on animal behaviour over time.

Several limitations should be noted. First, the study was conducted in a single small pasture (1.1 ha) with a single herd of Japanese Black cows, and the numbers of cows (22 and 24) were limited by herd size. Responses and rates of habituation may differ in larger pastures, with other breeds or physiological states, or under different sward conditions. Second, observers could not be blind to the device, because the device was visible in the videos. Third, the continuous trials used only the robot, so the comparison of habituation rates between devices relies on a previous study. Fourth, we did not measure physiological indicators such as heart rate, and stress responses may occur without overt behavioural signs (Ditmer et al., 2015). Finally, herding was operated manually; integrating the present findings with autonomous detection and decision-making is a key step towards robotic herding in routine grazing management.

In conclusion, a ground-level, dog-like robot elicited stronger avoidance and more successful herding than a drone, but responsiveness to the robot still declined over consecutive days, recovered partially after a pause and then declined again. These findings show how classical characteristics of habituation shape the responses of animals to robots under field conditions and identify habituation management as the central challenge for robotic herding.

## Supporting information

SAS code

## Data Availability

The data are openly available from Zenodo (Anzai et al., 2026; https://doi.org/10.5281/zenodo.20825305). The SAS code used for all analyses is provided as Supplementary Material.

## Declaration of Generative AI and AI-assisted Technologies in the Manuscript Preparation Process

During the preparation of this work, the authors used Claude (Anthropic) to edit the English language, to restructure the Introduction and Discussion, and to draft the SAS code for the statistical analyses. After using this tool, the authors reviewed and edited the content and the code as needed and take full responsibility for the content of the published article.

## Acknowledgments

We thank Ikuo Kobayashi, Genki Ishigaki, Koichiro Henmi and the staff of the Sumiyoshi Livestock Science Station, University of Miyazaki (Miyazaki, Japan), where this study was conducted, for animal management and field assistance, and Aya Nishiwaki for lending us the drone. We also thank Yosuke Goto, Minori Kan, Mizuki Tagashira, Moe Matsuhashi and Yui Kawano for field support. This work was supported by the Japan Society for the Promotion of Science (JSPS KAKENHI grant number 23K14047).

## Author Contributions

Hiroki Anzai: Conceptualization, Methodology, Formal analysis, Investigation, Data curation, Writing – original draft, Writing – review & editing, Visualization, Supervision, Project administration, Funding acquisition. Kirika Iwatani: Investigation, Data curation, Writing – original draft, Visualization. Sakuya Miyamoto: Investigation, Data curation, Writing – original draft, Visualization. Kanon Tamura: Investigation, Data curation, Writing – original draft. Honoka Miyagi: Investigation, Data curation, Writing – original draft.

## References

Anzai, H., & Kumaishi, M. (2023). Effects of continuous drone herding on behavioral response and spatial distribution of grazing cattle. Applied Animal Behaviour Science, 268, Article 106089. 10.1016/j.applanim.2023.106089

Anzai, H., & Sakurai, H. (2022). Preliminary study on the application of robotic herding to manipulation of grazing distribution: Behavioral response of cattle to herding by an unmanned vehicle and its manipulation performance. Applied Animal Behaviour Science, 256, Article 105751. 10.1016/j.applanim.2022.105751

Anzai, H., Iwatani, K., Miyamoto, S., Tamura, K., & Miyagi, H. (2026). *Data from: Animal-robot interaction in the field: Stronger responses and slower habituation of grazing cattle to a quadruped robot than to a drone* (Version 2) [Data set]. Zenodo. 10.5281/zenodo.20825305

Bolker, B. M., Brooks, M. E., Clark, C. J., Geange, S. W., Poulsen, J. R., Stevens, M. H. H., & White, J.-S. S. (2009). Generalized linear mixed models: A practical guide for ecology and evolution. Trends in Ecology & Evolution, 24(3), 127–135. 10.1016/j.tree.2008.10.008

Canteloup, C., Lee, J., Zimmermann, S., Alvino, M., Montenegro, M., Hutter, M., & van de Waal, E. (2024). *When monkeys meet an ANYmal robot in the wild* [Preprint]. bioRxiv. 10.1101/2024.08.13.607714

Distel, R. A., Soca, P. M., Demment, M. W., & Laca, E. A. (2004). Spatial–temporal arrangements of supplementation to modify selection of feeding sites by sheep. Applied Animal Behaviour Science, 89(1–2), 59–70. 10.1016/j.applanim.2004.04.006

Ditmer, M. A., Vincent, J. B., Werden, L. K., Tanner, J. C., Laske, T. G., Iaizzo, P. A., Garshelis, D. L., & Fieberg, J. R. (2015). Bears show a physiological but limited behavioral response to unmanned aerial vehicles. Current Biology, 25(17), 2278–2283. 10.1016/j.cub.2015.07.024

Doerfler, R. L., Lehermeier, C., Kliem, H., Möstl, E., & Bernhardt, H. (2016). Physiological and behavioral responses of dairy cattle to the introduction of robot scrapers. Frontiers in Veterinary Science, 3, Article 106. 10.3389/fvets.2016.00106

Herlin, A., Brunberg, E., Hultgren, J., Högberg, N., Rydberg, A., & Skarin, A. (2021). Animal welfare implications of digital tools for monitoring and management of cattle and sheep on pasture. Animals, 11(3), Article 829. 10.3390/ani11030829

King, A. J., Portugal, S. J., Strömbom, D., Mann, R. P., Carrillo, J. A., Kalise, D., de Croon, G., Barnett, H., Scerri, P., Groß, R., Chadwick, D. R., & Papadopoulou, M. (2023). Biologically inspired herding of animal groups by robots. Methods in Ecology and Evolution, 14(2), 478–486. 10.1111/2041-210X.14049

Kubinyi, E., Miklósi, Á., Kaplan, F., Gácsi, M., Topál, J., & Csányi, V. (2004). Social behaviour of dogs encountering AIBO, an animal-like robot in a neutral and in a feeding situation. Behavioural Processes, 65(3), 231–239. 10.1016/j.beproc.2003.10.003

Papadopoulou, M., Ball, M., Bartashevich, P., Burns, A. L. J., Chiara, V., Clark, M. A., Costelloe, B. R., Fele, M., French, F., Hauert, S., Heinrich, M. K., Herbert-Read, J. E., Hoitt, J., Ioannou, C. C., Landgraf, T., Matchette, S. R., Polverino, G., Sankey, D. W. E., Scott, D. M., … King, A. J. (2025). Active interactions between animals and technology: Biohybrid approaches for animal behaviour research. Animal Behaviour, 224, Article 123160. 10.1016/j.anbehav.2025.123160

Partan, S. R., Larco, C. P., & Owens, M. J. (2009). Wild tree squirrels respond with multisensory enhancement to conspecific robot alarm behaviour. Animal Behaviour, 77(5), 1127–1135. 10.1016/j.anbehav.2008.12.029

Rankin, C. H., Abrams, T., Barry, R. J., Bhatnagar, S., Clayton, D. F., Colombo, J., Coppola, G., Geyer, M. A., Glanzman, D. L., Marsland, S., McSweeney, F. K., Wilson, D. A., Wu, C.-F., & Thompson, R. F. (2009). Habituation revisited: An updated and revised description of the behavioral characteristics of habituation. Neurobiology of Learning and Memory, 92(2), 135–138. 10.1016/j.nlm.2008.09.012

Ravikanna, R., Cox, J., Heselden, J. R., Lloyd, R., Elias, A., & Hanheide, M. (2025). An exploratory study on the use of robot dogs in shepherding. In Proceedings of the 2025 20th ACM/IEEE International Conference on Human-Robot Interaction (HRI) (pp. 1820–1822). IEEE. 10.1109/HRI61500.2025.10974134

Réale, D., Reader, S. M., Sol, D., McDougall, P. T., & Dingemanse, N. J. (2007). Integrating animal temperament within ecology and evolution. Biological Reviews, 82(2), 291–318. 10.1111/j.1469-185X.2007.00010.x

Yaxley, K. J., Joiner, K. F., & Abbass, H. (2021). Drone approach parameters leading to lower stress sheep flocking and movement: Sky shepherding. Scientific Reports, 11, Article 7803. 10.1038/s41598-021-87453-y

