## Supplementary Material for "Grazing cattle respond more strongly to a quadruped robot than to a drone but habituate over consecutive days"

#### Table S1

Generalized linear mixed models of the responses of cows in the comparison trials

| Response | Effect | Estimate ± SE | df | *t* | *P* | OR (95% CI) |
|---|---|---|---|---|---|---|
| Move away | Intercept | −0.273 ± 0.239 | 21 | −1.15 | 0.265 | |
| | Device (robot) | 0.703 ± 0.260 | 242 | 2.70 | 0.007 | 2.02 (1.21–3.37) |
| | Sound (dog barking) | −0.041 ± 0.259 | 242 | −0.16 | 0.876 | 0.96 (0.58–1.60) |
| | Cow variance | 0.159 ± 0.147 | | | | |
| No response | Intercept | −1.235 ± 0.314 | 21 | −3.94 | <0.001 | |
| | Device (robot) | −1.843 ± 0.462 | 242 | −3.99 | <0.001 | 0.16 (0.06–0.39) |
| | Sound (dog barking) | −0.163 ± 0.404 | 242 | −0.40 | 0.688 | 0.85 (0.38–1.88) |
| | Cow variance | 0.213 ± 0.343 | | | | |
| Latency[a] | Threshold 0 s \| 1–9 s | −0.379 ± 0.274 | 19 | −1.38 | 0.183 | |
| | Threshold 1–9 s \| 10–19 s | 0.492 ± 0.276 | 19 | 1.78 | 0.091 | |
| | Threshold 10–19 s \| ≥20 s | 1.004 ± 0.283 | 19 | 3.54 | 0.002 | |
| | Device (robot) | 0.949 ± 0.257 | 242 | 3.70 | <0.001 | 2.58 (1.56–4.28) |
| | Sound (dog barking) | 0.188 ± 0.252 | 242 | 0.75 | 0.456 | 1.21 (0.73–1.99) |
| | Cow variance | 0.529 ± 0.338 | | | | |
| Flight | Intercept | 0.037 ± 0.220 | 21 | 0.17 | 0.868 | |
| | Device (robot) | 1.430 ± 0.286 | 242 | 5.00 | <0.001 | 4.18 (2.38–7.34) |
| | Sound (dog barking) | 0.123 ± 0.284 | 242 | 0.43 | 0.665 | 1.13 (0.65–1.98) |
| | Cow variance | 0.001 ± 0.105 | | | | |
| Herding success | Intercept | 0.499 ± 0.291 | 21 | 1.72 | 0.101 | |
| | Device (robot) | 1.459 ± 0.351 | 210 | 4.15 | <0.001 | 4.30 (2.15–8.59) |
| | Sound (dog barking) | −0.109 ± 0.333 | 210 | −0.33 | 0.744 | 0.90 (0.47–1.73) |
| | Cow variance | 0.365 ± 0.324 | | | | |

Binomial GLMMs with a logit link and cow identity as a random intercept (22 cows; $N = 266$ approach events, or 234 events with a known outcome for herding success). Reference levels: drone and air horn. Odds ratios (OR) compare the robot with the drone and dog barking with the air horn. Denominator degrees of freedom were obtained by the between–within method. [a] Cumulative logit GLMM for the four latency categories; an odds ratio above 1 indicates shorter latency.

### Table S2

Device × sound interaction terms in the comparison trials

| Response | Effect | Estimate ± SE | df | *t* | *P* |
|---|---|---|---|---|---|
| Move away | Device × sound | −0.073 ± 0.522 | 241 | −0.14 | 0.890 |
| No response | Device × sound | 0.828 ± 0.931 | 241 | 0.89 | 0.375 |
| Latency[a] | Device × sound | −1.080 ± 0.513 | 241 | −2.11 | 0.036 |
| | Robot vs drone, air horn | 1.409 ± 0.341 | 241 | 4.13 | <0.001 |
| | Robot vs drone, dog barking | 0.330 ± 0.389 | 241 | 0.85 | 0.397 |
| Flight | Device × sound | −0.209 ± 0.577 | 241 | −0.36 | 0.718 |
| Herding success | Device × sound | 1.379 ± 0.673 | 209 | 2.05 | 0.042 |
| | Robot vs drone, air horn | 0.833 ± 0.450 | 209 | 1.85 | 0.065 |
| | Robot vs drone, dog barking | 2.212 ± 0.528 | 209 | 4.19 | <0.001 |

Each interaction term was added to the model in Table S1. For herding success and latency, the effect of device (robot versus drone, log odds ratio) is also given separately for each sound. For herding success, odds ratios (95% CI) were 2.30 (0.95–5.58) with the air horn and 9.13 (3.23–25.84) with dog barking; for latency, they were 4.09 (2.09–8.02) with the air horn and 1.39 (0.65–2.99) with dog barking. [a] Cumulative logit GLMM.

**Responses of Calves to the Quadruped Robot**

Calves were not targeted but were present in the herding area throughout the continuous trials. Ten calves aged 74–176 days (mean ± SE = 135.7 ± 10.9 days) were present on Days 1–5, and 11 calves aged 68–204 days (mean ± SE = 152.0 ± 14.4 days) were present on Days 6–8. Calves could not always be identified individually in the videos, so their responses are summarized descriptively and were not analysed statistically. Responses were classified with the same methods as for cows.

We recorded 63 approach events involving calves across the eight trial days. Compared with cows (823 events), calves less often moved away (25.4% versus 43.0% of events) and more often showed interest (12.7% versus 5.1%), fear/startle/attack (7.9% versus 3.0%) and slight reactions (46.0% versus 32.8%). Calves more often responded with a latency of 0 s (71.4% versus 52.0%) and less often with a latency of 20 s or more (7.9% versus 22.4%), and they fled less often (47.6% versus 67.1%). Herding success was 85.7% for calves (18 of 21 approaches with a known outcome) and 81.4% for cows. These patterns suggest that calves were more likely to investigate the robot or to react promptly without leaving, consistent with greater exploration of novel objects by young animals.
